# Community sequencing on a natural experiment reveals little influence of host species and timing but a strong influence of compartment on the composition of root endophytes in three annual Brassicaceae

**DOI:** 10.1101/2020.01.24.918177

**Authors:** Jose G. Maciá-Vicente, Bora Nam, Marco Thines

## Abstract

The plant family Brassicaceae includes some of the most studied hosts of plant microbiomes, targeting microbial diversity, community assembly rules, and effects on host performance. Compared to bacteria, eukaryotes in the brassicaceous microbiome remain understudied, especially under natural settings. Here, we assessed the impact of host identity and age on the assembly of fungal and oomycete root communities, using DNA metabarcoding of roots and associated soil of three annual co-habiting Brassicaceae collected at two time points. Our results showed that fungal communities are more diverse and structured than those of oomycetes. In both cases, plant identity and sampling time had little influence on community variation, whereas root/soil compartment had a strong effect by exerting control on the entry of soil microorganisms into the roots. The enrichment in roots of specific fungi suggests a specialization towards the asymptomatic colonization of plant tissues, which could be relevant to host’s fitness and health.

## Introduction

Plants associate through their roots with microbial communities that are essential for their fitness, health, and response to environmental cues (Carrión et al., 2019; Durán et al., 2018; Kia et al., 2017). The elucidation of the mechanisms responsible for these associations, as well as the factors affecting them, provide new perspectives about how plant communities and ecosystems function (Klein et al., 2016; Wagg et al., 2019), and will likely enable the exploitation in agriculture of multiple aspects of plant-microbe symbioses (Duhamel and Vandenkoornhuyse, 2013; Wei and Jousset, 2017). Research focused on several members of the plant family Brassicaceae has spearheaded the advances in knowledge about the plant-associated microbiota (a.k.a. microbiomes) diversity and function (e.g., Almario et al., 2017; Bulgarelli et al., 2012; Dombrowski et al., 2016; Durán et al., 2018; Glynou et al., 2018a, 2018b; Keim et al., 2014; Lundberg et al., 2012; Wagner et al., 2016), both because of their economic importance and their inclusion of the most studied plant model species, *Arabidopsis thaliana* (L.) Heynh.

The microbiome of Brassicaceae species comprises multi-kingdom microbial communities dominated by bacteria, fungi, and oomycetes, which closely interact both with the host and among each other (Hassani et al., 2018). Most studies have focused on bacteria, as they may represent the largest fraction of the plant microbiome, have important effects on host performance, and have been recently proven instrumental in keeping at bay the detrimental effects caused by endophytic fungi and oomycetes on plant growth (Bulgarelli et al., 2012; Dombrowski et al., 2016; Durán et al., 2018; Lundberg et al., 2012; Wagner et al., 2016). However, the latter filamentous eukaryotes can also play important roles in microbiome function, encompassing likely keystone taxa that determine the structure of the microbial communities (Agler et al., 2016), well known pathogens that impact plant health and productivity, and mutualists able to assist hosts in the acquisition of nutrients (Almario et al., 2017; Hiruma et al., 2016).

Multiple studies have helped reveal the diversity and factors affecting the assembly of the eukaryotic microbiome of brassicaceous hosts, especially of fungi. However, a better resolution of the predominant fungi and oomycetes occurring in plants under natural conditions is still needed. Multiple studies engage in single sampling, have low spatial resolution with respect to the plant compartment investigated, or focus on common gardens with conditions different to those in natural habitats (but, for exceptions, see Glynou et al., 2018a, 2016; Thiergart et al., 2019). Besides, important factors that may affect the assembly of plant-associated microbial communities, such as plant identity and age or phenological status, have been addressed in bacteria (Dombrowski et al., 2016; Schlaeppi et al., 2014) but remain understudied in fungi and oomycetes. For example, the effect of brassicaceous host plant species in microbiome assembly has been mainly investigated across genotypes within a species (Agler et al., 2016; Glynou et al., 2016; Urbina et al., 2018), or using limited experimental settings that preclude generalization (Glynou et al., 2018b). The effect of time on the fungal and oomycete fraction of the microbiome has been studied across consecutive years (Thiergart et al., 2019), but finer resolution studies to evaluate the impact of host phenology are lacking.

Here, we address the impact of plant identity and age on the assembly of fungal and oomycete communities in root-associated soil and roots of annual Brassicaceae. We approach these questions by sampling at two time points — representing phenological growth stages of basal rosette and flowering — specimens of the brassicaceous plant species *A. thaliana, Cardamine hirsuta* L., and *Draba verna* L., co-habiting in a habitat undisturbed for decades. We targeted microbial communities at different rhizocompartments, ranging from bulk soil to the root endosphere, which have repeatedly shown to represent different degrees of host influence on microbial communities (so-called host-filtering effect) that increases toward the plant inner tissues (Almario et al., 2017; Bulgarelli et al., 2012; Lundberg et al., 2012; Martínez-Diz et al., 2019; Thiergart et al., 2019). We hypothesized that the effects of both host plant genotype and age on fungal and oomycete communities increases from soil to root compartments, owing to a host filtering effect on community composition that selects for specialized microorganisms with root-colonizing traits. Based on previous results on disturbed habitats, we expect a low influence of plant species in the assembly of fungal and oomycete communities (Glynou et al., 2018b), and a stronger effect of plant age due to normal seasonal changes in the soil pool of microbial species, coupled with metabolic changes in the host driven by phenology.

## Materials and Methods

### 2.1 Sample collection and process

The sampling site is located in west Germany (N 50.09844, E 8.54721, 114 m a.s.l.), consisting of an ancient gravel path in the north of the old castle of Frankfurt Hoechst, the sides of which were covered in moss that provided the habitat for various small annual plants. Populations of the Brassicaceae species *A. thaliana*, *C. hirsuta*, and *D. verna* co-habited at the site as interspersed stands, covering an area of approx. 50 m^2^. Samplings were performed at two time points: on February 20^th^, 2014, when plants were in a stage of basal rosette; and on April 20^th^, 2014, when plants presented a flowering stalk carrying fully developed flowers and siliques. At every sampling time, ten specimens of *A. thaliana*, and three specimens of each *C. hirsuta* and *D. verna* were collected by carefully uprooting the plants, taking care not to detach the soil particles adhered to roots. In addition, for each species, soil samples of approx. 10 cm^3^ were collected from just beneath each plant, and from approx. 20 cm apart in a spot free of flowering plants, using a small shovel that was rinsed and disinfected with isopropanol after each sample was taken. As the bulk soil was not expected to be very dissimilar across the site, only three bulk soil samples were taken for comparison with *A. thaliana* roots. Plant and soil samples were individually stored in zip-lock bags and brought to the laboratory in the same day, where they were kept at 4 C until processing on the next day.

For each plant individually, roots were separated from stems and soil adhering to the roots was removed by putting the roots into 50 ml reaction tubes half-filled with a sterile 0.01 % (v/v) Tween 20 solution and shaking for 10 min. After removal of the roots, the root-associated soil (rhizosphere soil) was pelleted by centrifugation and the supernatant removed before further processing. The washed roots were then separated into two halves, of which one was surface sterilized by incubating in 4 % (w/v) sodium hypochlorite solution with gentle shaking for about 20 s. Soil samples were homogenized and approximately 0.5 g of soil were mixed with the lysis buffer of the FastDNA™ SPIN Kit for Soil (MP Biomedicals, Solon, USA) and disrupted in a mixer mill (Retsch MM 200, Retsch GmbH, Haan, Germany) for 5 min at 25 Hz, using three iron beads with 3.5 mm per 2 ml reaction vial. After processing, samples represented five different root/soil compartments, each comprising 32 samples [(10 *A. thaliana* + 3 *C. hirsuta* + 3 *D. verna*) × sampling times]: bulk soil (soil collected apart from plant specimens), root zone soil (soil collected underneath plant specimens), rhizosphere (soil washed out of roots), root (non-sterilized roots), and endosphere (surface-sterilized roots).

### DNA extraction, amplification and sequencing

Total genomic DNA was extracted from root samples using the BioSprint 96 DNA Plant Kit (Qiagen GmbH, Hilden, Germany) on a KingFisher Flex 96 robotic system (Thermo Fisher Scientific, Waltham, MA, USA), and from soil samples with the FastDNA^™^ SPIN Kit for Soil, following the manufacturers’ instructions. DNA extracts were used directly, or after 10^-1^ dilution in molecular biology grade ddH_2_0 (VWR Chemicals, Darmstadt, Germany) for specific amplification and high throughput sequencing of the fungal rDNA internal transcribed spacer 1 (ITS1), and the oomycete cytochrome c oxidase subunit 2 (*cox2*) gene.

Amplification of the fungal ITS1 was done using primers ITS1F and ITS2, modified as in Smith and Peay (2014) to include the Illumina Nextera adapters, a linker sequence, and 12-bp error-correcting Golay barcodes (Table S4). PCR reactions were done in duplicate, in volumes of 25 μl, containing 1 μl of DNA template, 1× Phusion HF buffer (New England Biolabs GmbH, Schwalbach, Germany) with 1.5 mM MgCl_2_, 0.8 mg ml^-1^ bovine serum albumin (BSA, New England Biolabs GmbH), 0.2 mM of each dNTP (Bioline, Luckenwalde, Germany), 0.2 μM of each primer, and 0.5 units of Phusion Hot Start Flex DNA polymerase (New England Biolabs GmbH). Thermal cycles were carried out in a Mastercycler pro thermal cycler (Eppendorf, Hamburg, Germany) with the following program: 94 C for 1 min followed by 35 cycles of 94 C for 30 s, 52 C for 30 s, and 68 C for 30 s; and a final step of 68 C for 10 min. The amplicons were visualized in an electrophoresis agarose gel and volumes of individual PCR products between 6 and 12 μl were pooled based on band intensity (Duhamel et al., 2019). The DNA pool was purified with the Zymoclean TM Gel DNA Recovery Kit (Zymo Research, Freiburg, Germany), quantified with a Qubit Fluorometer (Thermo Fisher Scientific), and paired-end sequenced by Eurofins Genomics GmbH (Ebersberg, Germany) with the Illumina MiSeq platform, using the MiSeq Reagent Kit v3 (Illumina Inc., San Diego, CA, USA).

For amplification of the oomycete *cox2* gene, primers Cox2-F and Cox2-RC4 (Choi et al., 2015; Hudspeth et al., 2000) were used (Table S4). PCR reactions were done as described above, but with the polymerase MangoTaq (Bioline, Luckenwalde, Germany), using the following thermal conditions: 94 C for 4 min followed by 36 cycles of 94 C for 40 s, 53 °C for 20 s, and 72 C for 60 s; and a final step of 72 C for 4 min. Library preparation and sequencing with the MiSeq platform was performed by LGC Genomics GmbH (Berlin, Germany).

The sequence data generated in this study has been deposited in the NCBI Sequence Read Archive under BioProject number PRJNA593383.

### Sequence processing

Sequence reads from the fungal ITS1 region and the oomycete *cox2* gene were processed using the DADA2 pipeline (Callahan et al., 2016) for quality filtering, dereplication, removal of chimeric sequences, and grouping of reads into exact amplicon sequence variants (ASVs; Callahan et al., 2017). In the case of the ITS1, the process included the merging of paired forward and reverse reads after the dereplication step. The paired-end merging was omitted in the case of *cox2* sequences given the frequent lack of overlapping regions between forward and reverse reads, owing to the length of the amplicon. In this case, only forward reads were processed due to their overall higher quality respect to the reverse reads. The code used to process both paired-end and single direction sequences is available online at https://github.com/jgmv/MiSeq_process.

Taxonomic annotation of ASV sequences was done with the Naïve Bayesian Classifier tool (Wang et al., 2007) available in mothur v1.39.5 (Schloss et al., 2009). For the fungal ITS1, annotation was achieved by comparing sequences against the UNITE database of fungal ITS sequences (Kõljalg et al., 2005). Because no similar database is available for the oomycete *cox2* gene, in this case we used an *ad hoc* built reference sequence data set with all oomycete cox2 sequences available in NCBI GenBank as of July 26^th^, 2019.

### Statistical analyses

All statistical analyses were carried out in R v3.6.1 (R Core Team, 2019). Both the ITS1 and *cox2* data sets were processed independently, but using the same procedures. First, ASVs represented globally by less than five reads were discarded, as well as nine samples from the *cox2* dataset that contained no reads. Sampling coverage was assessed by the construction of rarefaction curves per sample, using functions in R package vegan v2.5-4 (Oksanen et al., 2019). These showed marked differences in read abundances and ASV richness, although in most cases saturation in the ASV accumulations indicated appropriate coverage of samples’ richness (Fig. S1). We relied on a mixture model to normalize reads and account for differences in library size and biological variability, using the variance stabilization method with package DESeq v1.35.1 (Anders and Huber, 2010; Love et al., 2014). This method has shown to outperform rarefying based-normalization because it does not require samples or species with few reads to be discarded and does not decrease the statistical power in analyses (McMurdie and Holmes, 2014).

Diversity and community structure analyses relied on functions in package vegan. ASV richness and diversity, based on the Shannon diversity index (H’) and represented as effective species numbers (*e*^H’^; Jost, 2006), were calculated per sample using the non-normalized datasets. Significant differences in reads abundance, richness, and diversity across plant species and compartments were assessed by means of the Kruskal-Wallis rank sum tests. In the case of comparisons of between sampling dates, paired Wilcoxon rank sum tests were used. Kruskal-Wallis tests with Bonferroni adjustment of *P* values to account for multiple comparisons were applied to assess significant variations in the abundance of individual taxa across soil/root compartments. Pearson’s correlation coefficient (*r*) was used to assess relationships between read numbers and richness or diversity values per sample.

To investigate differences in community composition across plant species, soil/root compartment, and sampling date, we calculated Bray-Curtis dissimilarities among samples in the normalized datasets, and visualized them using unconstrained principal coordinates analysis (PCoA) ordinations. To further assess the contribution of variables to community variation, constrained distance-based redundancy analysis (db-RDA) was applied, using plant species, soil/root compartment, and sampling date as constraining factors.

We investigated the specific association of fungal and oomycete ASVs with rhizospheric and plant compartments by assessing their enrichment or depletion respect to bulk soil, following a differential abundance analysis adapted from that described in Edwards et al. (2015). For this, data from compartments bulk soil and root zone soil were combined, and the fold change abundance of individual ASVs in rhizosphere, root, and endosphere compartments vs. soil were calculated based on the coefficients of fitted generalized linear models (GLM) of abundance data. The non-normalized data set with read counts was used to build the models based on a negative binomial regression, with read numbers across samples accounted for by including them as a model parameter (i.e. *abundance* ~ *reads* + *compartment*). Differential abundance was calculated based on the GLM coefficients for the rhizosphere, root, and endosphere compartments, representing fold change respect to soil, and tested by the likelihood ratio test with an adjusted *P* value cutoff of 0.01. Positive coefficient values indicate enrichment of and ASV’s abundance in a compartment respect to its abundance in soil, whereas negative values indicate depletion (Edwards et al., 2015).

All the data sets and the bash and R code used for the data pre-processing and statistical analyses has been made available online at https://github.com/jgmv/Brassicaceae_fungal_and_oomycete_root_microbiome.

## Results

A total of 7,567,288 sequence reads representing 4,289 fungal ASVs were retained in the fungal ITS dataset after quality filtering and discarding rare ASVs. For the oomycete *cox2* dataset, 4,105,321 reads representing 951 ASVs were kept. In the case of *cox2* data, nine samples from the root and endosphere compartments of *A. thaliana* and *D. verna* were dropped due to absence of reads (Table 1). Despite attempts to normalize the quantity of amplified DNA prior to MiSeq sequencing, there was a strong effect of plant compartment on the number of ITS reads per sample, with roots and endosphere showing significantly lower read numbers (Kruskal-Wallis test, H = 50.8, df = 4, *P* < 0.01; Table 1, Fig. S2) than soil samples. This effect was not significant in the case of *cox2* reads, neither had plant species a strong effect on either ITS and *cox2* read numbers (Table 1, Fig. S2). Likewise, plant compartment, but not plant species, affected fungal (H = 98.4, df = 4, *P* < 0.01) and oomycete (H = 80.5, df = 4, *P* < 0.01) ASV richness. Date of sampling had a significant effect on the number of reads (paired Wilcoxon test, V = 1752, *P* < 0.001) and ASV richness (V = 1687.5, *P* < 0.001) per sample in the fungal ITS dataset, but not in the oomycete dataset (Fig. S2).

**Table 1.** Distribution of numbers of reads and amplicon sequence variants (ASVs) richness across fungal and oomycete samples.

|  |  | Fungi |  |  |  |  |  | Oomycota |  |  |  |  |  |
| --- | --- | --- | --- | --- | --- | --- | --- | --- | --- | --- | --- | --- | --- |
|  |  | February |  |  | April |  |  | February |  |  | April |  |  |
|  |  | n | Reads / plant<sup>1</sup> | Richness | n | Reads / plant | Richness | n | Reads / plant | Richness | n | Reads / plant | Richness |
| *A. thaliana* | Bare soil | 3 | 57121 ± 9202 | 686 | 3 | 97170 ± 54434 | 759 | 3 | 41594 ± 21817 | 73 | 3 | 10954 ± 7898 | 59 |
|  | Root soil | 3 | 69249 ± 1304 | 743 | 3 | 69060 ± 28821 | 847 | 3 | 28300 ± 10051 | 64 | 3 | 48440 ± 35566 | 58 |
|  | Rhizosphere | 10 | 35093 ± 22781 | 999 | 10 | 94449 ± 78228 | 1522 | 10 | 33911 ± 31632 | 192 | 10 | 34644 ± 26163 | 214 |
|  | Root | 10 | 21189 ± 29870 | 214 | 10 | 130423 ± 176687 | 313 | 10 | 33190 ± 33411 | 44 | 9 | 56005 ± 37747 | 88 |
|  | Endosphere | 10 | 14630 ± 43466 | 89 | 10 | 37609 ± 99336 | 128 | 8 | 50724 ± 92048 | 20 | 8 | 33853 ± 53642 | 20 |
| *C. hirsuta* | Bare soil | 3 | 36908 ± 25218 | 501 | 3 | 73449 ± 19219 | 870 | 3 | 38093 ± 24432 | 67 | 3 | 7636 ± 5203 | 46 |
|  | Root soil | 3 | 62464 ± 14017 | 737 | 3 | 113586 ± 79217 | 789 | 3 | 25880 ± 8770 | 56 | 3 | 22933 ± 23163 | 50 |
|  | Rhizosphere | 3 | 55227 ± 3139 | 694 | 3 | 134748 ± 101309 | 762 | 3 | 21413 ± 17630 | 70 | 3 | 19186 ± 10021 | 69 |
|  | Root | 3 | 44161 ± 50344 | 74 | 3 | 190322 ± 133137 | 126 | 3 | 80482 ± 138248 | 16 | 3 | 50206 ± 47608 | 13 |
|  | Endosphere | 3 | 523 ± 282 | 31 | 3 | 4041 ± 6329 | 43 | 3 | 39416 ± 68215 | 7 | 3 | 4180 ± 5795 | 6 |
| *D. verna* | Bare soil | 3 | 41529 ± 18834 | 606 | 3 | 60902 ± 4129 | 810 | 3 | 25264 ± 23975 | 67 | 3 | 46958 ± 54960 | 61 |
|  | Root soil | 3 | 52975 ± 17889 | 573 | 3 | 84638 ± 11754 | 939 | 3 | 31846 ± 22576 | 51 | 3 | 15610 ± 18422 | 57 |
|  | Rhizosphere | 3 | 69498 ± 2484 | 710 | 3 | 70873 ± 26902 | 870 | 3 | 19708 ± 18113 | 62 | 3 | 20748 ± 20390 | 70 |
|  | Root | 3 | 4472 ± 4044 | 40 | 3 | 9982 ± 13726 | 59 | 3 | 6103 ± 10295 | 7 | 3 | 30661 ± 39981 | 20 |
|  | Endosphere | 3 | 439 ± 121 | 16 | 3 | 7780 ± 11554 | 30 | 1 | 285 ± 0 | 1 | 1 | 93 ± 0 | 1 |

<sup>1</sup> Values represent the mean number of reads per plant ± standard deviation.

Shannon diversity was less affected by read abundances than ASV richness (*r* = 0.01, *P* = 0.9 vs. *r* = 0.26, *P* = 0.003 in the ITS dataset; *r* = 0.12, *P* = 0.2 vs. *r* = 0.17, *P* = 0.06 in the *cox2* dataset), and hence was further used to evaluate diversity patterns across factors (Fig. 1). Again, the soil/root compartment significantly affected diversity in both fungal (H = 81.1, df = 4, *P* < 0.01) and oomycete (H = 81.2, df = 4, *P* < 0.01) communities, most markedly by clearly lower numbers of effective ASVs in the root and endosphere as compared to soil compartments (Fig. 1). Plant species and sampling date did not show a relationship with fungal or oomycete diversity, although samples collected in April appeared to have larger variation in diversity values than those collected in February (Fig. 1).

**Figure 1.**
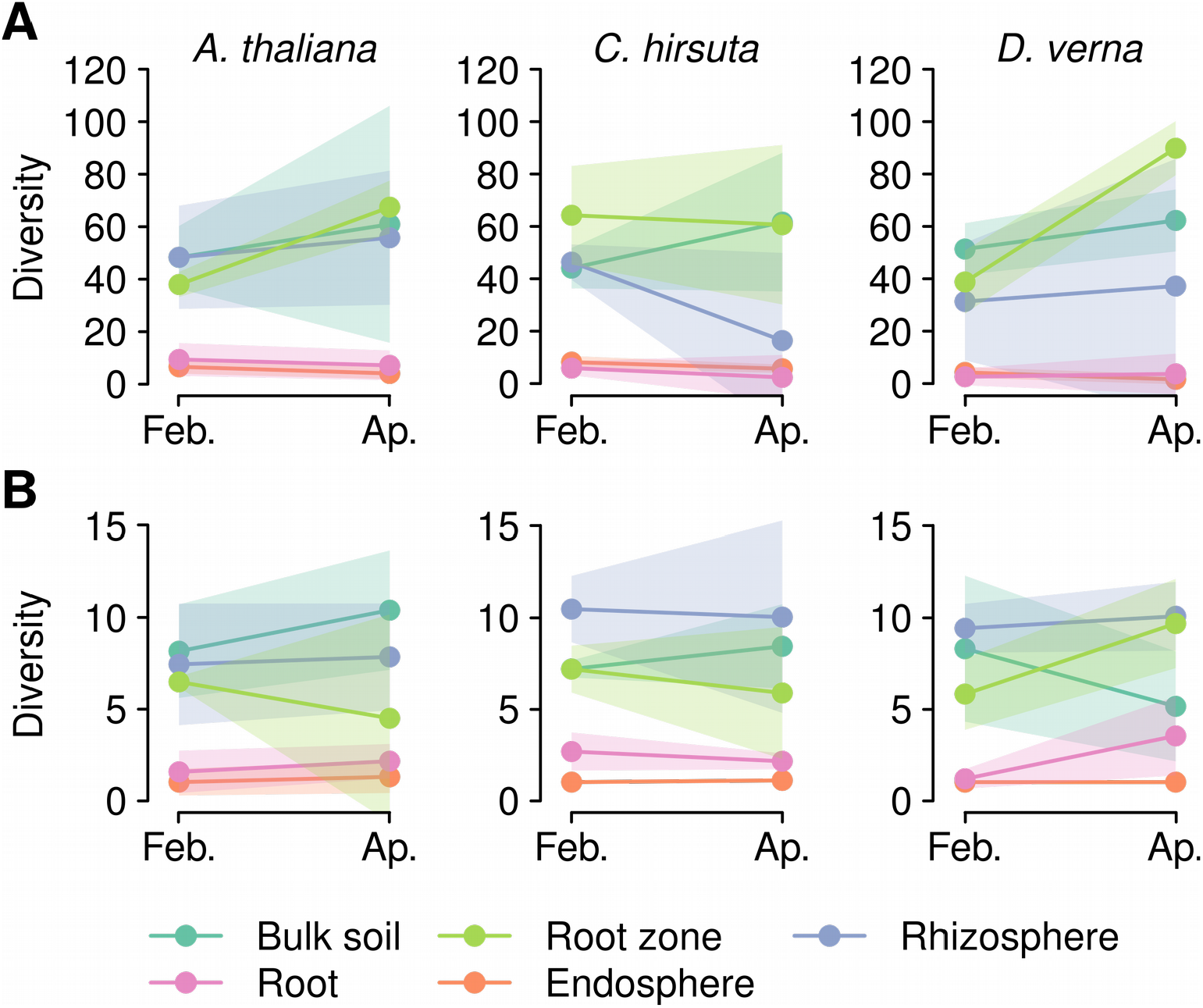
Diversity of fungal (**A**) and oomycete (**B**) communities across plant species (different plots), sampling times (x axis: **Feb**., February; **Ap**., April), and soil/root compartments (point and line colors). Diversity is expressed as effective ASV numbers, calculated based on the Shannon diversity index. Points represent the mean, and polygons the standard deviation of diversity values.

The ordination analysis of fungal and oomycete community structures showed similar patterns to those of diversity, with samples from soil and root compartments forming two main, separate clusters (Fig. 2A,C). db-RDA ordination of fungal communities explained a significant 26.2 % of overall community variation (pseudo-*F*_7,124_ = 6.3, *P* = 0.001). Of this variation, the largest fraction was related to differences in soil/root compartment (Fig. 2B), with all soil compartments forming a compact cluster well separated from root and endosphere samples, which also tended to cluster separately from each other (Fig. 2A). A similar pattern was found for oomycetes, although in this case the db-RDA only explained an 8.5 % of community variation (pseudo-*F*_7,115_ = 1.5, *P* = 0.001) and, even though compartment was again the most explanatory factor (Fig. 2E), there was a lesser definition of the db-RDA clusters (Fig. 2D). Separate db-RDA ordinations were calculated for the different soil/root compartments to assess the effects of plant species and sampling date on different fractions of the microbial communities (Figs. 2C,F). These showed in all cases that plant species explained a larger fraction of community variation than sampling date, although this was relatively low and did not change markedly across compartments.

**Figure 2.**
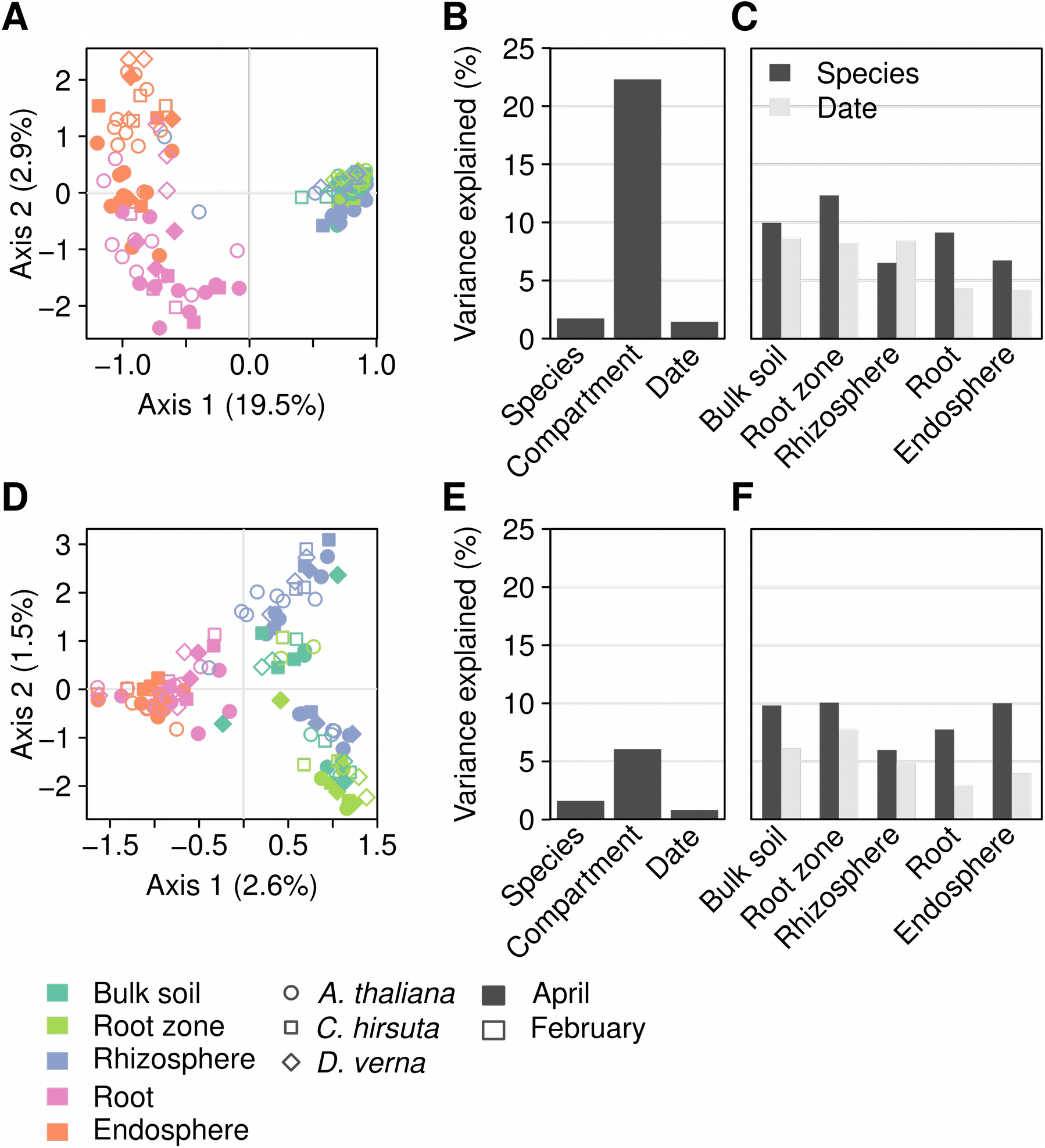
Effect of host identity, date of sampling, and soil/root compartment on the structure of fungal (**A** – **C**) and oomycete (**D** – **F**) communities. Plots in **A** and **C** show db-RDA ordinations based on Bray-Curtis dissimilarities, constrained by host, date, and compartment. **B** and **D** show the variance explained in db-RDAs by each of the three factors, whereas **C** and **F** show the variance explained by host species and date in subsets of data for each soil/root compartment.

Most fungal ASVs belonged in the Ascomycota (48.5 %), followed by the Basidiomycota (20 %), the Glomeromycota (2.2 %), and the Mortierellomycota (2.2 %); and were classified into 86 orders, plus 17 and 3 taxa unclassified or with uncertain classification (*Incertae sedis*) at the order level (Table S1). The most abundant orders showed a significant variation in their relative abundance across soil/root compartments (Table S2), which was consistent across plant species (Fig. 3A). This was most evident by an increase in the representation of ASVs in the Pleosporales (H = 91.7, df = 4, *P_adj_*, < 0.001), Helotiales (H = 85.3, df = 4, *P_adj_* < 0.001), Olpidiales (H = 32.6, df = 4, *P_adj_* < 0.001), and Cantharellales (H = 88, df = 4, *P_adj_* < 0.001) in the root compartments; and by the opposite pattern in the Chaetothyriales (H = 103.6, df = 4, *P_adj_* < 0.001) and Mortierellales (H = 99.4, df = 4, *P_adj_* < 0.001), as well as in the unclassified Fungi (H = 99.3, df = 4, *P_adj_* < 0.001). In oomycetes, most ASVs remained unclassified at all taxonomic levels (Fig. 3A, Table S3). Those that could be assigned to an order were distributed across 5 orders, with the Phythiales comprising the largest number of ASVs (34.5 %), followed by far by the Peronosporales (5.3 %) and the Saprolegniales (3 %). In this case, no evident patterns in the distribution of the orders abundances across soil/root compartments was found.

**Figure 3.**
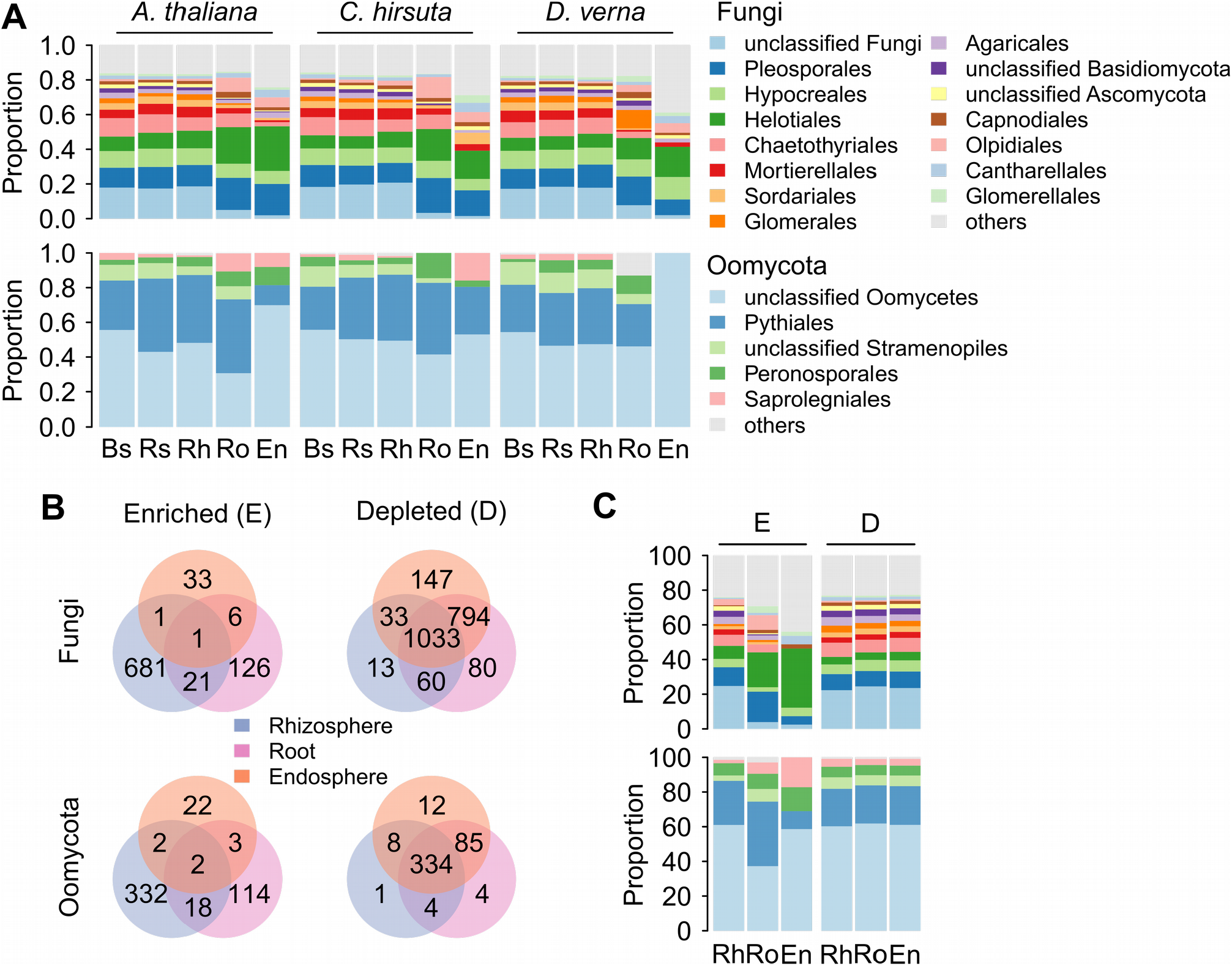
Taxonomic structure of fungal and oomycete communities and variation of ASV’s abundance across soil/root compartments. Bar plots in **A** show the distribution of fungal (top) and oomycetes (bottom) orders across host species and soil/root compartments. Venn diagrams in **B** show the numbers of differentially enriched or depleted ASVs in rhizocompartments as compared to bulk soil. **C** shows the proportion of enriched (**E**) and depleted (**D**) ASVs in different rhizocompartments within the main fungal and oomycete orders. Abbreviations: **Bs**, bulk soil; **Rs**, root zone soil; **Rh**, rhizosphere; **Ro**, root; **En**, endosphere.

The differential abundance analyses of ASVs showed that fungi were much more depleted than enriched in rhizosphere and root compartments respect to soil, with 2160 depleted vs. 869 enriched ASVs, whereas oomycetes appeared to be enriched or depleted at similar rates (Fig. 3B; Table S3). In both fungi and oomycetes, enrichment of ASVs occurred mostly in the rhizosphere. On the contrary, most ASVs were depleted in all compartments, although others were exclusively depleted in both root compartments (Fig. 3B). In all cases, several ASVs appeared to be exclusively enriched or depleted at specific compartments (Fig. 3B). In fungi, the root-enriched ASVs belonged mostly to the Helotiales, which increased their representation respect to soil both in the root surface and in the endosphere (Fig. 3C; Table 2). They were followed by the Pleosporales and Pezizales (solely represented by the genus *Tuber*), which became enriched in the outer and inner root, respectively (Fig. 3C; Table 2). In oomycetes, the major enrichment in roots included *Pythium rostratifingens, Phytophthora citricola*, and *Saprolegnia ferax*, as representatives of the Pythiales, Peronosporales, and Saprolegniales (Fig. 3C; Table 2). The ASVs depleted in rhizosphere and plant compartments belonged to multiple fungal and oomycete orders (Fig. 3C), somewhat mirroring the overall distribution of orders found for both groups (Fig. 3A).

**Table 2.** Fungal and oomycete amplicon sequence variants (ASVs) enriched in the endosphere compartment.

|  |  |  | Host<sup>2</sup> |  |  |
| --- | --- | --- | --- | --- | --- |
|  | ASV | Compartment<sup>1</sup> | *A.*<br>*thaliana* | *C.*<br>*hirsuta* | *D.*<br>*verna* |
| Fungi |  |  |  |  |  |
| *Calycina* sp. (Helotiales) | FU00009 | e | + | + | + |
| *Tuber* sp. (Pezizales) | FU00160 | e,r | + | + | + |
| *Kabatiella* sp. (Dothideales) | FU00212 | e,r | + | - | + |
| *Paraphoma* sp. (Pleosporales) | FU00277 | e | + | + | + |
| unclassified Helotiales | FU00433 | e,r | + | - | + |
| unclassified Helotiales | FU00607 | e | + | - | - |
| unclassified Coniochaetales | FU00766 | e,r | + | + | + |
| unclassified Sebaciniales | FU00802 | e | + | - | - |
| *Athelia* sp. (Atheliales) | FU00851 | e,r | + | + | + |
| *Apiotrichum xylopi*<br>(Trichosporonales) | FU00893 | e | + | - | + |
| *Aspergillus* sp. (Eurotiales) | FU00930 | e,r | + | + | + |
| unclassified Sordariomycetes | FU00953 | e | + | + | + |
| *Stemphylium* sp. (Pleosporales) | FU00961 | e,r | + | - | - |
| unclassified Dothideomycetes | FU00966 | e | + | + | + |
| *Calycina ellisii* (Helotiales) | FU01001 | e | + | - | - |
| unclassified Ceratobasidiaceae<br>(Cantharellales) | FU01006 | e | + | + | - |
| unclassified Sebaciniales | FU01030 | e | + | - | - |
| *Dactylonectria* sp. (Hypocreales) | FU01089 | e | + | + | + |
| unclassified Hydnodontaceae<br>(Trechisporales) | FU01292 | e | + | - | - |
| unclassified Rutstroemiaceae<br>(Helotiales) | FU01388 | e | - | + | - |
| *Tetracladium marchalianum*<br>(Helotiales) | FU01433 | e | - | - | + |
| unclassified Nectriaceae<br>(Hypocreales) | FU01515 | e | - | - | + |
| unclassified Helotiales | FU01572 | e | - | + | - |
| unclassified Thelebolaceae<br>(Thelebolales) | FU01626 | e | + | - | - |
| *Tuber borchii* (Pezizales) | FU01638 | e | - | + | - |
| unclassified Helotiales | FU01696 | e | + | - | - |
| unclassified Helotiales | FU01775 | e | + | - | - |
| *Plectosphaerella cucumerina*<br>(Glomerellales) | FU01776 | e | - | + | - |
| *Tetracladium marchalianum*<br>(Helotiales) | FU01777 | e | - | - | + |
| *Articulospora* sp. (Helotiales) | FU01797 | e | + | - | - |
| *Tuber borchii* (Pezizales) | FU01828 | e | + | - | - |
| *Tuber oligospermum* (Pezizales) | FU01829 | e | - | - | + |
| *Rhizodermea* sp. (Helotiales) | FU01870 | e | - | - | + |
| unclassified Fungi | FU01943 | e | - | - | + |
| *Thanatephorus cucumeris*<br>(Cantharellales) | FU01967 | e | - | + | - |
| *Tetracladium marchalianum*<br>(Helotiales) | FU02008 | e | + | - | - |
| *Tuber oligospermum* (Pezizales) | FU02049 | e | + | - | - |
| *Apiotrichum gracile*<br>(Trichosporonales) | FU02050 | e | + | - | - |
| *Tetracladium marchalianum*<br>(Helotiales) | FU02071 | e | + | - | - |
| *Tuber oligospermum* (Pezizales) | FU02240 | e | - | - | + |
| unclassified Capnodiales | FU02499 | e | + | - | - |
| **Oomycota** |  |  |  |  |  |
| unclassified Oomycetes | OO00004 | e,r | + | + | + |
| unclassified Oomycetes | OO00005 | e | + | + | - |
| unclassified Oomycetes | OO00024 | e,r | + | - | - |
| *Pythium rostratifyingens* (Pythiales) | OO00028 | e | + | - | + |
| unclassified Saprolegniaceae<br>(Saprolegniales) | OO00047 | e | + | + | - |
| *Pythium rostratifyingens* (Pythiales) | OO00048 | e,r | + | - | + |
| unclassified Oomycetes | OO00063 | e | + | - | - |
| unclassified Oomycetes | OO00066 | e | + | - | - |
| unclassified Oomycetes | OO00067 | e | - | + | - |
| *Phytophthora citricola*<br>(Peronosporales) | OO00071 | e,r | + | - | - |
| unclassified Oomycetes | OO00075 | e | + | - | - |
| unclassified Oomycetes | OO00088 | e | + | - | - |
| *Saprolegnia ferax* (Saprolegniales) | OO00117 | e | - | + | - |
| unclassified Oomycetes | OO00122 | e | + | - | - |
| *Saprolegnia ferax* (Saprolegniales) | OO00142 | e | + | - | - |
| unclassified Oomycetes | OO00154 | e | + | + | + |
| unclassified Oomycetes | OO00167 | e | + | - | - |
| *Phytophthora citricola*<br>(Peronosporales) | OO00195 | e | + | - | - |
| *Saprolegnia ferax* (Saprolegniales) | OO00317 | e | + | - | - |
| unclassified Oomycetes | OO00478 | e | + | - | - |
| unclassified Oomycetes | OO00485 | e | - | - | + |
| unclassified Oomycetes | OO00679 | e | + | - | - |
| unclassified Oomycetes | OO00689 | e,r | + | + | - |
| unclassified Pythiales | OO00755 | e | + | - | - |
| unclassified Oomycetes | OO00793 | e | + | - | - |
| *Saprolegnia ferax* (Saprolegniales) | OO00801 | e | - | + | - |
| unclassified Peronosporales | OO00836 | e | + | - | - |
| unclassified Peronosporaceae<br>(Peronosporales) | OO00916 | e | - | + | - |
| unclassified Oomycetes | OO00920 | e | + | - | - |

<sup>1</sup> Whether the ASVs was detected in non-sterilized roots.Plant compartment where the ASVs were enriched: r, root; e, endosphere.
<sup>2</sup> Hosts in which the ASVs were detected.

## Discussion

The fungal and oomycete communities in roots of three brassicaceous plant species and their associated soil were barely affected by host identity and a time of sampling spanning two months. Instead, the major driver of community differences was the soil/root compartment, which determined differences in community diversity and structure between the microbiota developing in soil and different parts of the roots. These results suggest that the interface between soil and roots exerts a strong barrier to root colonization, but also that those fungi and oomycetes able to surpass the root boundaries are generalist groups without strong host preferences.

The remarkable differences between soil and root-associated fungal and oomycete communities described here have been already reported for multiple microbial groups and plant lineages. Studies on several plant species have found compartments, ranging from root-associated soil to the root endosphere, to be the strongest determinants of community variation, following patterns similar to those found here, both in fungi (Almario et al., 2017; Coleman-Derr et al., 2016; Durán et al., 2018; Martínez-Diz et al., 2019; Thiergart et al., 2019) and oomycetes (Durán et al., 2018; Thiergart et al., 2019), but also in other microbial groups like protists (Sapp et al., 2018) and bacteria (Bulgarelli et al., 2012; Coleman-Derr et al., 2016; Durán et al., 2018; Edwards et al., 2015; Lundberg et al., 2012; Thiergart et al., 2019). Altogether, there is strong evidence that the root surface exerts a control on the entry of soil microorganisms into the root tissues. This control has been proposed to depend on two main processes: an enrichment from soil towards the root tissues of soil microbes with root-colonizing traits, starting already at a certain distance from the rhizoplane probably in response to root exudates; and an exclusion of the largest fraction of soil microorganisms, mostly taking place at the rhizoplane level (Edwards et al., 2015; Heijden and Schlaeppi, 2015). This model of microbial recruitment, proposed for bacteria, was only partly met by our results, and it varied between fungi and oomycetes. Here, the blockade of microbial entry into the roots was already observed away from the root surface, since most ASVs were jointly depleted from the rhizosphere toward the inner root tissues, and appeared to be unspecific because most orders decreased proportionally to their overall abundance. Bacteria that respond to root exudates by locally increasing their populations and activating antimicrobial functions have been shown to play a role in the suppression of fungal root pathogens at the rhizosphere level (Chapelle et al., 2016; Leveau et al., 2010), a process that could be responsible for unspecifically inhibiting growth of other fungi and oomycetes (Durán et al., 2018). This could also explain the lack of an important distance enrichment of ASVs in the rhizosphere, as has been observed for bacteria (Edwards et al., 2015). In addition, few or no enriched ASVs were shared among the rhizosphere, root, and endosphere compartments, indicating that each of these likely represent specialized niches for different taxa.

Plant species did not affect considerably community structure, neither from fungi nor from oomycetes. Neither this effect increased in root compartments respect to the surrounding soil, where the influence of species-specific chemistry, defense responses, or microbial recognition, would expectably lead the processes of microbial enrichment and depletion described above. However, this finding is in line with previous reports of a limited host specificity of root-associated microbes (Bulgarelli et al., 2012; Colin et al., 2017; Dombrowski et al., 2016; Glynou et al., 2018b; Schlaeppi et al., 2014; Thiergart et al., 2019; Wagner et al., 2016; Zarraonaindia et al., 2015). This is at least true in related plant species, such as those investigated in our study, because phylogenetically distant hosts harbor clearly distinguishable microbial communities (U’Ren et al., 2019) likely due to major differences in cellular physiology and chemistry. The ability of endophytes to colonize multiple hosts appears to be frequent among non-pathogenic symbionts (Põlme et al., 2018), in stark contrast to the high host specialization found throughout the pathogenic lifestyle (Thines, 2019). The loose host preferences of endophytes may enable them to persist in and disseminate across habitats with diverse communities of potential plant hosts.

As with host identity, the sampling time did not affect significantly the structure of microbial communities in soil or roots. Particularly in roots, the temporal effect would have been ascribed to two processes: the normal seasonal variation in soil microbial communities, which would have been detectable in plant-free soil samples; and the physiological changes in root tissues upon phenological stages, which would likely have strongest effects in root compartments. Changes in soil microbiota have been shown to occur over seasons, although they are smaller than those triggered by other ecological factors, such as habitat type or spatial distance, and they are more evident over longer periods of time than those considered in this study (He et al., 2017). Regarding phenotype-driven modifications of the plant microbiome, Dombrowski et al, (2016) showed that non-flowering wild types of the brassiaceous species *Arabis alpina* assembled bacterial root communities indistinguishable from those of flowering mutants, and concluded that the root microbiota is robust to changes following its established in earlier stages of plant development. Our results reinforce this conclusion, expanding it to eukaryotic members of the root microbiome. A broad phylogenetic diversity of fungi were found associated with roots of the three Brassicaceae species studied, reflecting the taxonomic fungal diversity in the surrounding soil. As this study, previous works have repeatedly shown a consistent dominance of root fungal endophytic communities by members of the Pleosporales, Hypocreales, and Helotiales (Bonito et al., 2014; Glynou et al., 2018a, 2018b; Keim et al., 2014; Knapp et al., 2012), which likely occupy complementary niches within roots (Kia et al., 2019). However, our new data suggest that prevalence of hypocrealean fungi might depend on their soil abundance, as they were not importantly enriched in root tissues. The Helotiales were particularly enriched in the root and endosphere compartments, in line with previous studies and the known association of this order with plant roots, where they might be involved in assisting host nutrition, as has been shown in some instances (Almario et al., 2017; Johnston et al., 2019). In contrast to fungi, enrichment patterns in oomycetes were less clear, and involved groups of well-known root pathogens like *Pythium* and *Phytophtora*. In this case, most ASVs remained unclassified. Even though the *cox2* gene has been shown adequate for species identification and phylogenetics within the oomycetes (Choi et al., 2015), the low diversity of *cox2* sequences represented in public repositories presently hinders its application as barcoding marker.

Our sample processing yielded an overall lower sequencing depth in the root than in soil samples, and that was particularly low in the endosphere. These differences in sampling coverage across compartments may have influenced to some extent the results on diversity and community structure, reducing resolution in ASV detection. These differences likely reflect the low amount of fungal and oomycete DNA, and hence biomass, within plant tissues. This problem when amplifying endophytic DNA has been overcome elsewhere by the application of nested PCR prior to sequencing (Eusemann et al., 2016; Unterseher et al., 2016), although we opted to avoid this procedure due to potential risks of distorting relative abundances and community structures detected (Yu et al., 2015). Nevertheless, the diversity and community patterns detected are consistent with those obtained in similar studies, thus indicating a valid biological signal in our results.

In conclusion, our results show that fungi assemble more diverse and complex communities than oomycetes in plant roots and their associated soil. However, microbial communities of both groups are similarly structured across rhizocompartments, and relatively lowly affected by plant identity and phenology of annual Brassicaceae sharing the same habitat. The enrichment of specific lineages in roots, particularly in fungi, suggests a level of specialization towards the asymptomatic colonization of below-ground plant tissues, hinting symbiotic lifestyles which could be relevant to host’s fitness and health. The ecological meaning of these interactions, however, remains unknown and warrants further study.

## Acknowledgements

This study was supported by LOEWE (Landes-Offensive zur Entwicklung Wissenschaftlic-ökonimischer Exzellenz) of the state of Hesse and was conducted within the framework of the Cluster for Integrative Fungal Research (IPF). J.G.M.-V. acknowledges support from the German Research Foundation under grant MA7171/1-1.

## Author Contribution

M.T. designed the study. B.N. and M.T. collected the samples. B.N. processed the samples and prepared the DNA extracts. J.G.M.-V. and B.N. prepared the libraries for Illumina MiSeq sequencing. M.T. and J.G.M.-V. contributed materials and reagents. J.G.M.-V. analyzed the data and prepared a first draft of the manuscript, with input from B.N. and M.T. All authors contributed to the final version of the manuscript.

## Supplementary information

**Table S1.** Taxonomic classification of fungal and oomycete ASVs.

**Table S2.** Variation across soil/root compartments in the abundance of fungal and oomycete orders.

**Table S3.** Significant (*P* < 0.01) differential abundances of fungal and oomycete ASVs across compartments.

**Table S4.** Primers used in this study.

**Figure S1.** Rarefaction curves of ASVs accumulation with sequence reads, for the fungal ITS (**A**) and the oomycete *cox2* gene (**B**) datasets.

**Figure S2.** Reads (**A**) and ASVs richness (**B**) values obtained by Illumina MiSeq sequencing across the factors considered in this study. Box-and-whisker plots summarize the distribution of each measurement (median, interquartile range, and range) per factor, and points show individual values per sample.

