## Supplementary figures for "Community sequencing on a natural experiment reveals little influence of host species and timing but a strong influence of compartment on the composition of root endophytes in three annual Brassicaceae"

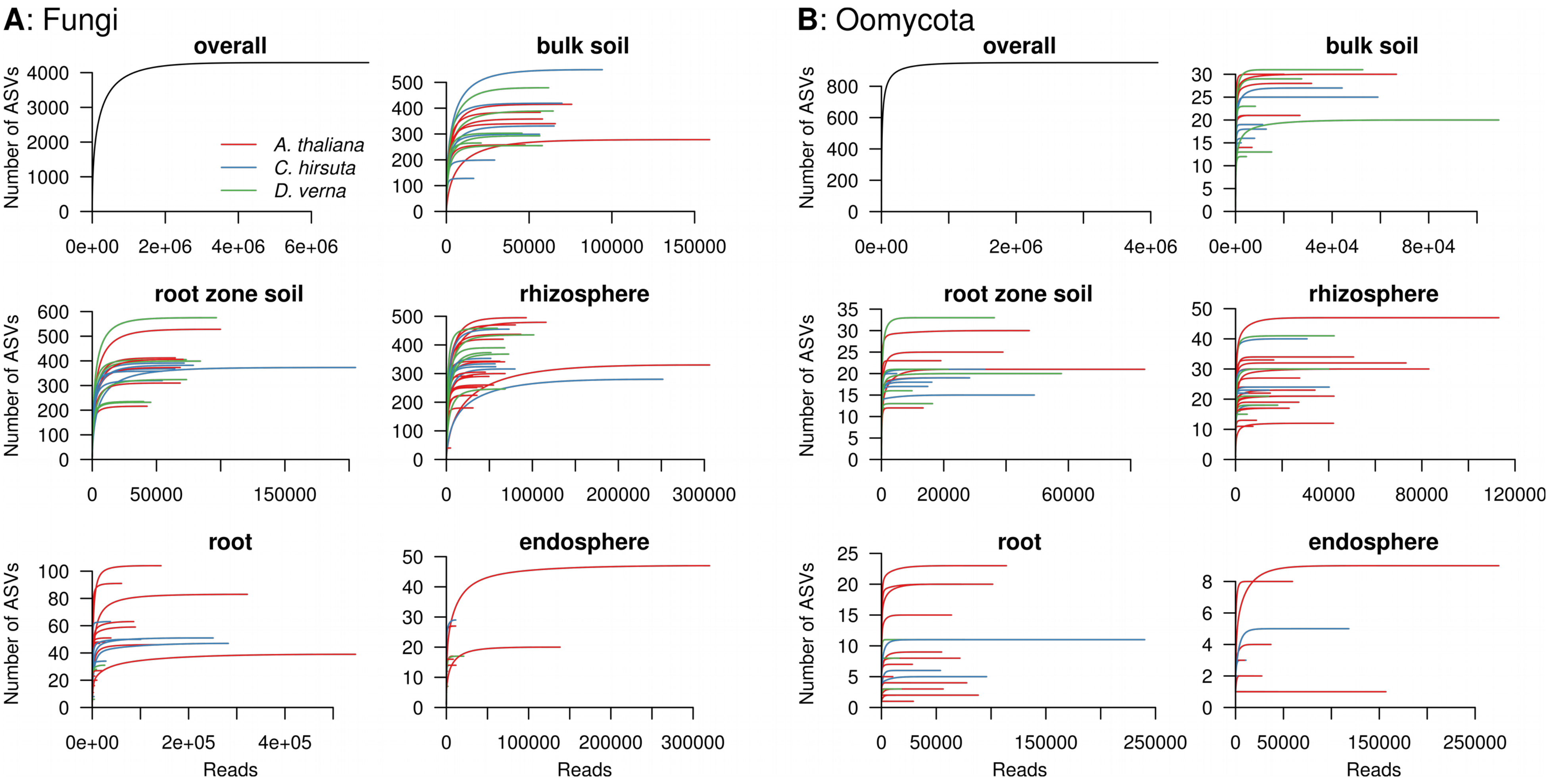


**Figure S1.** Rarefaction curves of ASVs accumulation with sequence reads, for the fungal ITS (**A**) and the oomycete *cox2* gene (**B**) datasets.

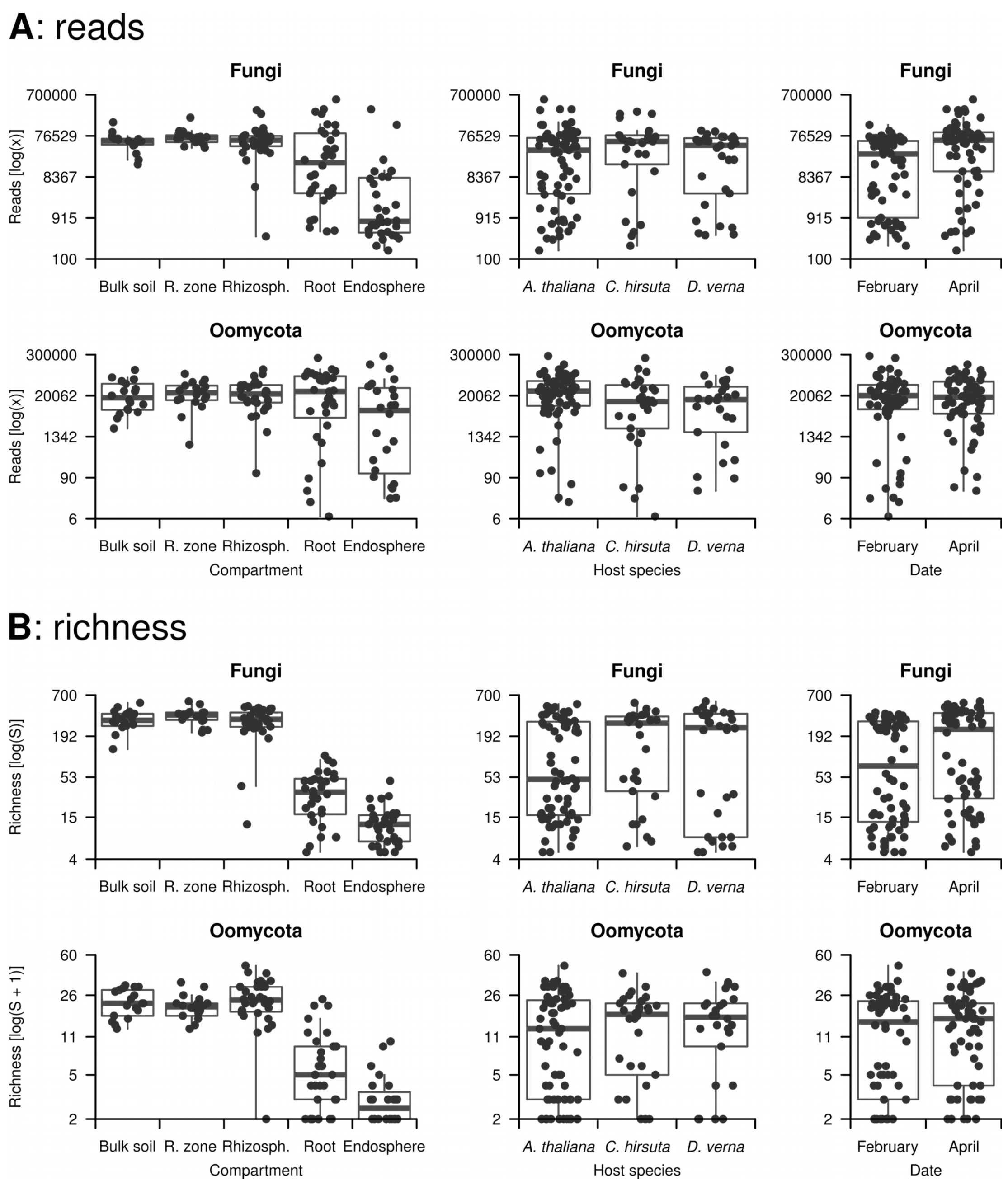


**Figure S2.** Reads (**A**) and ASVs richness (**B**) values obtained by Illumina MiSeq sequencing across the factors considered in this study. Box-and-whisker plots summarize the distribution of each measurement (median, interquartile range, and range) per factor, and points show individual values per sample.
